# 4-methylumbelliferone attenuates amyloid pathology and learning deficits in the APP/PS1 mouse model

**DOI:** 10.64898/2026.05.21.726929

**Authors:** Matthew Amontree, James O’Leary, Pauline Wonnenberg, Mateo Nelson, Katherine Conant

**Author notes:** Corresponding author: Katherine Conant, Georgetown University School of Medicine, 3970 Reservoir Road, Washington DC 20007; Electronic mail.

## Abstract

4-Methylumbelliferone (4-MU) inhibits hyaluronic acid (HA) synthesis and is currently approved in Europe for biliary spasm. 4-MU administration reduces perineuronal nets (PNNs), and enzymatic degradation of PNNs in mouse models of Alzheimer’s disease (AD) attenuates memory impairment. Although 4-MU has therapeutic efficacy in rodent models of fibrosis and cancer, it has not been examined in an Alzheimer’s model. Here, we evaluated the impact of long-term 4-MU treatment in the APP/PS1 amyloid mouse model. From three months of age, mice were on either a vehicle or 4-MU–supplemented diet for 70 days or 52 weeks. Short and long-term 4-MU treatment decreased the soluble parenchymal Aβ_1-42_/Aβ_1-40_ ratio. Reductions in insoluble amyloid plaque were observed following 52 weeks of treatment. Extended 4-MU administration also reduced PNN intensity and ameliorated spatial memory deficits in APP/PS1 mice. These findings provide support for targeting brain extracellular matrix (ECM) as a therapeutic strategy for AD.

## Introduction

AD is the most common dementia with >42 million people living with this disease globally and this is expected to triple in prevalence to 131 million people by 2050 ^1^. Hallmarks of AD include forebrain cholinergic neuronal loss, extracellular amyloid plaques, intracellular hyper-phosphorylated tau aggregates, vascular leakiness, and chronic neuroinflammation ^2^. Current treatment options include acetyl-cholinesterase inhibitors and anti-Aβ immunotherapies. Cholinesterase inhibitors delay cognitive loss, but efficacy diminishes with disease progression as cholinergic synapses are degraded ^3^. Anti-Aβ immunotherapies target the underlying pathology and delay cognitive loss but have yet shown the ability to halt disease progression. Given that AD neuropathology can precede cognitive impairment by >20 years ^4^, there is an urgent need for improved diagnostics and therapeutics.

The ECM of the central nervous system is particularly enriched in chondroitin sulfate proteoglycans (CSPGs), which are distributed throughout the diffuse ECM and concentrated within specialized matrix structures such as perineuronal nets (PNNs) ^5^. PNNs form a lattice-like sheath surrounding the soma and proximal dendrites of select neurons. In the hippocampus and cortex, these structures are most prominently associated with parvalbumin (PV)–expressing GABAergic interneurons, where they play a key role in neuronal maturation and the maintenance of fast-spiking activity ^6^. PNN assembly is initiated by the host neuron through the secretion of HA from membrane-bound HA synthases. Members of the lectican family, including aggrecan, neurocan, versican, and brevican, bind to the HA backbone via their N-terminal globular domains, an interaction stabilized by hyaluronan and proteoglycan link proteins (HAPLNs). The resulting complexes are further cross-linked at their C-terminal domains by tenascins, generating the characteristic net-like architecture of PNNs. Functionally, PNNs contribute to synaptic stabilization, and their emergence coincides with the closure of developmental critical periods ^7,8^. By reinforcing established synaptic connections while constraining the formation of new ones, PNNs act as regulators of neuroplasticity and cognitive flexibility ^9,10^. Consistent with this role, enzymatic degradation of PNNs using chondroitinase ABC (ChABC) has been shown in multiple studies to reduce Aβ pathology and improve behavioral outcomes *in vivo* ^11–13^. Moreover, Yang and colleagues reported that ChABC treatment enhances astrocytic autophagy–lysosome pathway activity, providing a potential mechanistic link between ECM remodeling and amyloid clearance ^13^.

4-MU is a coumarin derivative that inhibits HA synthesis through multiple mechanisms. Classically, 4-MU suppresses HA production by depleting intracellular pools of UDP-glucuronic acid, a critical substrate for HA biosynthesis ^14^. Because 4-MU is a competitive substrate for UDP-glucuronyl transferases (UGTs), relatively high concentrations are required to overcome endogenous glucuronic acid levels ^15^. In addition to this substrate-depletion mechanism, several studies have demonstrated that 4-MU downregulates transcription of HA synthase enzymes, providing an additional route by which HA synthesis is reduced ^14,16,17^. Notably, UDP-glucuronic acid also serves as a precursor for chondroitin sulfate (CS) synthesis, and prior work has shown that 4-MU treatment decreases C6S–positive PNN labeling *in vivo* ^18,19^. However, it remains unclear whether this effect reflects a direct reduction in CS biosynthesis or occurs secondarily through destabilization of the HA backbone that anchors PNNs. Beyond its effects on ECM components, 4-MU has been reported to enhance insulin sensitivity and lower blood glucose levels in diabetic mouse models ^20–22^. Chronic administration is well tolerated in rodents and has been associated with a substantial extension of lifespan, approximately 32 weeks compared with vehicle-treated controls ^23^. Long-term 4-MU treatment has also been shown to increase extracellular space volume (ECS) while reducing cortical and hippocampal fractional anisotropy in rats, indicating broader effects on brain tissue microstructure ^24^.

In our study we investigated the therapeutic efficacy of 4-MU treatment in the APP/PS1 amyloid mouse model. We sought to determine if 4-MU demonstrated the ability to reduce amyloid pathology and attenuate learning and memory deficits. Furthermore, we characterized the effects of chronic 4-MU treatment on the CNS ECM architecture. Also, we assessed the effects of 4-MU in primary cultured glia to determine if it directly reduces CS levels.

## Materials & Methods

### Mice

The National Institute of Health (NIH) ethical guidelines were followed, and protocols were approved by Georgetown University’s Institutional Animal care and Use Committee (IACUC). Wild-type (WT) C57BL/6J (000664) and hemizygous APP/PS1 on a C57BL/6J background (034832) mice were purchased from Jackson laboratories. Two female WT mice were paired with one APP/PS1 male mouse for breeding. Mice were ear punched at 2 months of age for genotyping to determine for presence of APP/PS1 transgenes. The following primers were used to determine genotype: 5’-GTGTGATCCATTCCATCAGC-3’, 5’-GGATCTCTGAGGGGTCCAGT-3’, and 5’-ATGGTAGAGTAAGCGAGAACACG-3’. Until 3 months of age, mice were given a standard chow diet by Georgetown University’s Division of Comparative Medicine (DCM). From 3 months of age until completion of the study, mice were given either a medicated diet containing 5% 4-MU (wt/wt) with chocolate flavoring or a non-medicated chocolate flavored diet to serve as the vehicle. 4-MU was purchased from Sigma-Aldrich (M1381) and the diets were produced by Bio-Serv (Flemington, NJ). Following completion of the study, mice were euthanized. Mice were anesthetized with isoflurane and decapitated with scissors. The brain was rinsed with cold PBS and hemi-sected with a razor. One hemi-section was placed into 4% paraformaldehyde for 24 hours, rinsed with PBS, and placed in 30% sucrose solution at 4° C until sliced on a microtome. The other hemi-section was micro dissected to collect the hippocampus and pre-frontal cortex. The tissues were sonicated in RIPA lysis buffer containing protease and phosphatase inhibitor (ThermoFisher, 78440), centrifuged at 13,300 x g for 20 minutes, and the supernatant was collected and stored at -80° C. The total protein in tissue lysates was determined using Pierce BCA assay (ThermoFisher, 23225).

### Rotarod

The rotarod assay is used to assess motor ability in rodents. The rotarod assessment was conducted across three consecutive days, consisting of two training sessions followed by a testing session. Mice were first allowed to acclimate to the behavioral testing room for 30 minutes. During each training day, animals completed three trials in which they remained on the rotarod while it rotated at a constant speed of 10 RPM for 5 minutes. On the testing day, mice again completed three trials, but the rod accelerated from 4 to 40 RPM over a 5-minute period, and performance values were averaged across trials. The primary outcome measure was latency to fall, defined as the time until the animal disengaged from the rotating rod. The apparatus was disinfected with 70% ethanol between individual mouse sessions.

### Barnes Maze

The Barnes maze was used to evaluate spatial learning and memory in mice. The protocol consisted of four days of acquisition training followed by a probe trial conducted three days after the final training session. The circular maze contained 20 evenly spaced holes arranged into four quadrants. Nineteen openings functioned as decoys that did not provide sufficient shelter, whereas one opening led to an escape box that allowed the animal to fully hide from the brightly lit exposed platform. A schematic representation of the apparatus is shown in **figure 1**. Distinct visual cues composed of different shapes (Ex: triangles) and patterns were positioned on each surrounding wall to provide spatial orientation. During training, mice were placed at the center of the maze, and the time required to locate and enter the escape box (escape latency) was recorded. Animals that failed to enter the escape within the 180-second cutoff were gently guided into the escape box and assigned an escape latency of 180 seconds. Each mouse completed three trials per day with approximately 1 hour between trials. For the probe trial, the escape box was removed and replaced with a decoy identical to the other 19 openings. Mice were allowed to explore the maze for 180 seconds starting from the center. The primary outcome measures were the latency to first enter the target zone corresponding to the former escape location, and total time spent within that zone. A reversal learning phase began the day after the initial probe trial and followed the same structure (four training days and one probe day). In this phase, the escape box was relocated to a different quadrant, requiring the animals to suppress the previously learned location and acquire the new one. The maze was cleaned with 70% ethanol between each mouse to eliminate odor cues.

**Figure 1:**
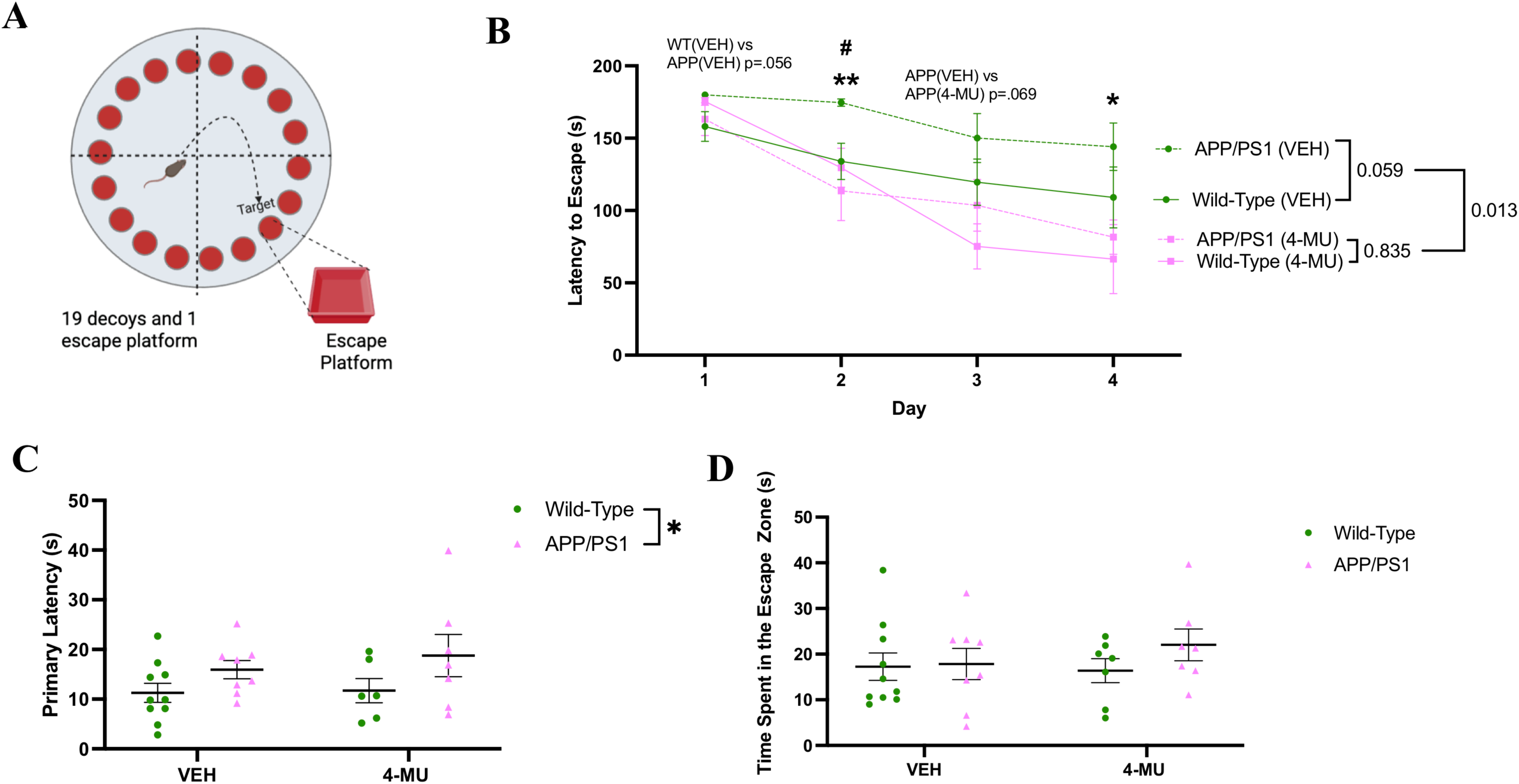
Chronic 4-MU treatment improves learning on the Barnes maze assay. **(A)** Graphical schematic of Barnes maze. The schematic was created in BioRender. **(B)** Average escape latency of the training component of the test. 2-way repeated measures ANOVA with Šidák’s multiple comparisons test. #Comparisons between WT (VEH) and APP/PS1 (VEH), #p<0.05; *Comparisons between APP/PS1 (VEH) and APP/PS1 (4-MU) **p<0.01, *p<0.05 **(C)** Primary latency of mice on the probe day. One outlier removed in WT (4-MU) group via ROUT’s test (Q=1%). 2-way ANOVA. **(D)** Total time spent in the escape proximity zone on probe day. 2-way ANOVA. n=7-10 per group, mean ± SEM, *p<0.05.

### Western Blot

Western blot analysis was conducted using mouse brain tissue and cell lysates. Equivalent protein concentrations were loaded for brain and cell lysates. For detection of CSPGs, samples were first treated with chondroitinase ABC (ChABC; Sigma-Aldrich, C3667) for 30 minutes at 37° C. Samples were then combined with Laemmli buffer containing 5% β-mercaptoethanol, heated at 85° C for 10 minutes, cooled on ice, separated by SDS-PAGE gel electrophoresis, and transferred to nitrocellulose membranes (Bio-Rad, 1704159). Membranes were stained with Ponceau S (Cell Signaling Technology, 59803S), blocked in 5% non-fat milk with 0.05% tween-20 in TBS (Bio-Rad, 1706404), and incubated overnight at 4° C with primary antibodies diluted in 3% BSA with 0.05% tween-20 in TBS on an orbital shaker. The following day, membranes were washed three times (10 minutes each) in TBS containing 0.05% tween-20 and incubated for 1 hour at room temperature with HRP-conjugated secondary antibodies (1:5000) diluted in 5% non-fat milk with 0.05% tween-20 in TBS. Membranes were subsequently washed three additional times in 0.05% tween-20 in TBS and twice in TBS to remove residual detergent. Chemiluminescent signal was developed using either West Pico PLUS substrate (ThermoFisher, 34579) or SuperSignal™ West Femto substrate (ThermoFisher, 34095). Ponceau S staining was used to assess transfer quality and for normalization during densitometric analysis. For re-probing with subsequent primary antibodies, membranes were incubated in Restore Western blot stripping buffer (ThermoFisher, 21059) for 30 minutes at room temperature on an orbital shaker. Stripped membranes were washed five times in TBS with 0.05% tween-20 before application of the next primary antibody. Primary antibodies used included neurocan N-terminal (1:2000, R&D Systems, AF5800), neurocan C-terminal (1:2000, Invitrogen, PA5-79718), brevican (1:4000, BD Biosciences, 610894), versican (1:1000, Abcam, EPR12277), aggrecan (1:1000, Sigma-Aldrich, ab1031), HAPLN1 (1:10,000, NSJ Bioreagents, RQ5347), HAPLN2 (1:2000, Neuromab, N364/10), chondroitin-6-sulfate (1:100, Amsbio, 270433-CS), 4-chondroitin-4-sulfate (1:200,000, Millipore, MAB2030), and APP (1:1000, clone 6E10, Novus, NBP2-62566). Secondary antibodies were goat anti-rabbit IgG-HRP (1:5000, Invitrogen, 31460), goat anti-mouse IgG-HRP (1:5000, Invitrogen, 31430), and donkey anti-sheep IgG-HRP (1:5000, Invitrogen, A16047).

### Immunofluorescence (IF)

Mouse brains were sliced at 40 μm on a microtome and were collected as free-floating sections in cryoprotectant solution (30% ethylene glycol, 30% sucrose, and phosphate buffer). Four dorsal hippocampal sections were selected for each mouse. Sections were first washed three times in TBS (10 minutes each) before antigen retrieval. For antigen retrieval, slices were steamed for 3 minutes in 10 mM citrate buffer containing 0.05% tween-20 (pH= 6.0) using a rice cooker. After cooling in TBS for 5 minutes, tissues were permeabilized with 0.3% triton X-100 in TBS for 10 minutes, then blocked for 1 hour in 10% goat serum with 0.1% triton X-100 in TBS. Slices were subsequently incubated overnight at 4° C on an orbital shaker in primary antibody solution consisting of 1% goat serum and 0.1% triton X-100 in TBS. The following day, sections were washed three times for 10 minutes in TBS containing 0.1% triton X-100 and then incubated for 2 hours at room temperature with secondary antibodies and/or Wisteria Floribunda Agglutinin (WFA) diluted in 1% goat serum with 0.1% triton X-100 in TBS. Tissues were washed three additional times in 0.1% triton X-100, rinsed once in TBS for 5 minutes, stained with DAPI for 10 minutes, and given a final 5 minute TBS wash. Slices were mounted using Vectashield vibrance mounting medium (VWR, H-1800), allowed to dry overnight at room temperature, and stored in light-protected boxes at 4° C until imaging. Primary antibodies included aggrecan (1:1000, Sigma-Aldrich, ab1031), parvalbumin (PV; 1:1000, Millipore, P3088), and APP (1:500, clone 6E10, Novus, NBP2-62566). Fluorophore-conjugated WFA (VWR, FL-1351-2) was used at 1:500. Secondary antibodies were goat anti-rabbit IgG (H+L) highly cross-adsorbed Alexa Fluor™ 647 (1:1000, Invitrogen, A-21245), goat anti-mouse IgG (H+L) highly cross-adsorbed Alexa Fluor™ Plus 555 (1:1000, Invitrogen, A-21422), and goat anti-rabbit IgG (H+L) highly cross-adsorbed Alexa Fluor™ 488 (1:1000, Invitrogen, A-11008).

### ELISA

Hyaluronan levels were measured using the DuoSet ELISA from R&D Systems (DY3614) in mouse brain tissue lysates and mouse glial conditioned media. Aβ_1-42_ (KMB3481) and Aβ_1-40_ (KMB3441) were measured with ELISAs from ThermoFisher. Values were normalized to the total protein concentration of each sample.

### Primary Mixed Glial Cultures

Primary mixed glial cultures were prepared from postnatal day 0–2 C57BL/6J mouse pups obtained from breeder pairs purchased from Jackson Laboratory (000664). Animals were euthanized by decapitation, brains were rapidly removed, cortical tissue was isolated by microdissection, and the meninges were carefully removed using forceps. Cortices were minced in Hank’s balanced salt solution (HBSS), transferred to complete medium (Minimum Essential Medium [MEM] supplemented with 1 mM L-glutamine, 1 mM sodium pyruvate, 0.6% D-glucose, 100 μg/mL penicillin/streptomycin, 4% fetal bovine serum, and 6% horse serum), centrifuged at 1,000 RPM for 5 minutes, and resuspended in growth media. Cells were plated into T-75 culture flasks and maintained in a humidified incubator at 37 °C with 5% CO . On day 21 *in vitro*, flasks were shaken at 200 RPM for 3 hours at 37° C to dislodge the superficial microglial layer. The adherent mixed glial layer was washed with PBS, incubated with trypsin-EDTA for 5–10 minutes, and enzymatic activity was quenched with excess media. The cell suspension was centrifuged at 200 × g for 10 minutes to pellet cells, and the supernatant was discarded. Cells were then resuspended in MEM containing 10% FBS, 1 mM L-glutamine, and 100 μg/mL penicillin/streptomycin and plated in 12-well plates. Before experimental treatments, serum-containing medium was removed, cultures were rinsed once with PBS, and serum-free MEM (1 mM L-glutamine and 100 μg/mL penicillin/streptomycin) was applied for 2 hours. This serum deprivation step was used to suppress astrocyte proliferation and growth.

## Results

### 52 weeks of 4-MU treatment improves spatial memory and subsequent reversal learning in APP/PS1 mice

Spatial memory deficits are noted in humans with AD ^25^ and in transgenic AD mouse models ^26^. Thus, we assessed if 52 weeks of 4-MU treatment could improve the performance of 15-month-old APP/PS1 on the Barnes maze which assesses spatial learning and memory. 4-MU administration significantly decreased escape latencies during the training phase relative to vehicle treatment (p=0.013), indicating improved task acquisition **(Figure 1)**. Across genotypes, APP/PS1 mice tended to exhibit longer escape latencies than WT mice, although this difference did not reach statistical significance (p=0.059). Within the APP/PS1 cohort, mice receiving 4-MU performed significantly better than vehicle-treated counterparts on training days 2 and 4 (p<0.05), with a trend observed on day 3 (p=0.069). Long-term spatial memory was evaluated using a probe trial conducted three days after the final acquisition session, during which the escape box was removed and replaced with a decoy. Memory performance was quantified by the latency to first enter the former escape location (primary latency) and the cumulative time spent within that zone. No effects of 4-MU treatment were detected for either outcome measure. In contrast, APP/PS1 mice displayed a significantly prolonged primary latency compared with WT mice (p=0.0378), consistent with impaired spatial memory retention. Reversal learning, which may reflect cognitive flexibility, was assessed one day later by relocating the escape box to a novel quadrant, requiring suppression of the previously learned location and formation of a new spatial association. During reversal acquisition, mice treated with 4-MU showed significantly faster learning than vehicle-treated animals regardless of genotype (p=0.002) **(Figure 2)**. A subsequent reversal probe trial performed three days later revealed no significant treatment-related differences in either primary latency or time spent in the new target zone (all p-values> 0.05).

**Figure 2:**
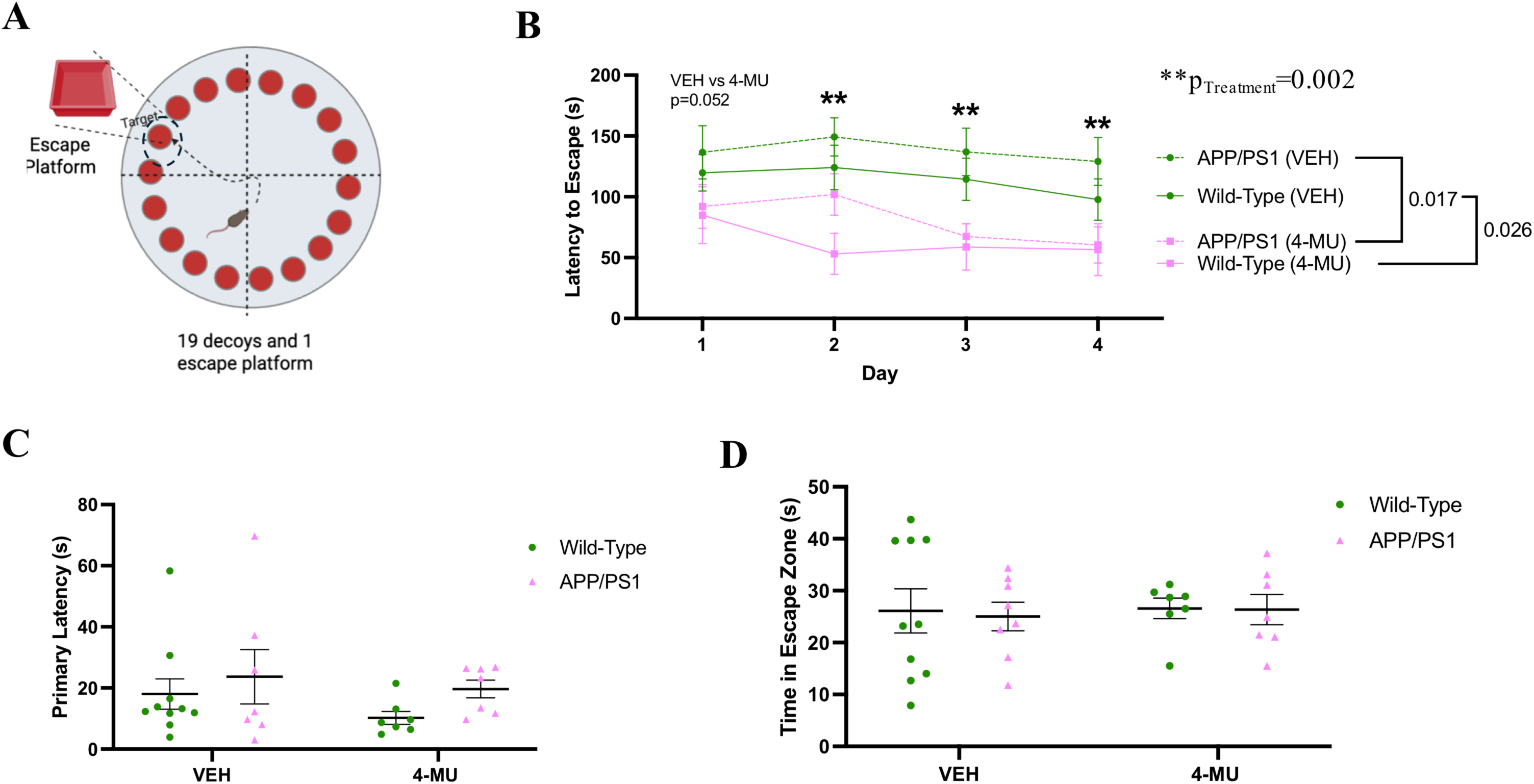
Chronic 4-MU treatment improves reversal learning. **(A)** Graphical schematic of reversal Barnes maze phase. The schematic was created in BioRender. **(B)** Average escape latency of the training phase of the reversal Barnes maze assay. 2-way repeated measures ANOVA with Šidák’s multiple comparisons test. **p<0.01, *p<0.05. **(C)** Primary latency of mice on the reversal probe day. 2-way ANOVA. **(D)** Total time spent in the escape proximity zone on reversal probe day. 2-way ANOVA. n=7-10 per group, mean ± SEM.

### Chronic 4-MU treatment modulates the hippocampal ECM architecture

We assessed the impact of 70 days and 52 weeks of 4-MU treatment on ECM components and PNN fluorescent intensity in WT and APP/PS1 mice using western blot, IF, and ELISA. IF was employed to quantify PNNs, as their components are also present in the diffuse ECM, whereas western blotting allowed discrimination between intact proteins and their cleavage fragments. Both 70-day and 52-week 4-MU treatments selectively reduced hippocampal chondoroitin-6-sulfate (C6S) levels **(Figure 3)**, while chondroitin-4-sulfate (C4S) levels remained unchanged. No genotype-dependent differences in C6S or C4S levels were observed in either experiment. In the 70-day study, two high molecular weight (>250 kDa) C6S bands were resolved and analyzed separately. 4-MU treatment significantly decreased levels of band 1 [F(1,18)= 5.776, p= 0.0272] and band 2 [F(1,19)= 17.26, p= 0.00005] relative to vehicle. In the 52-week experiment, the two high molecular weight bands could not be distinguished and were quantified together, revealing a significant reduction in total C6S levels following 4-MU treatment [F(1,23)= 7.245, p = 0.0130]. No significant changes in hippocampal C4S levels were detected at either time point (data not shown). Importantly, we found that hippocampal HA levels were reduced with both 70 days and 52 weeks of 4-MU treatment **(Figure 3)**. 70 days of 4-MU treatment significantly reduced hippocampal HA levels [F(1,19)= 12.85, p= 0.002]. Also, we observed this reduction in HA levels with 52 weeks of 4-MU treatment [F(1, 27)= 7.923, p= 0.009]. Unexpectedly, APP/PS1 mice at both 5.5 months of age [F(1,19)= 20.51, p=0.0002] and at 15 months of age [F(1,27)= 5.164, p= 0.0313] showed a significant reduction in hippocampal HA levels compared to WT littermates.

**Figure 3:**
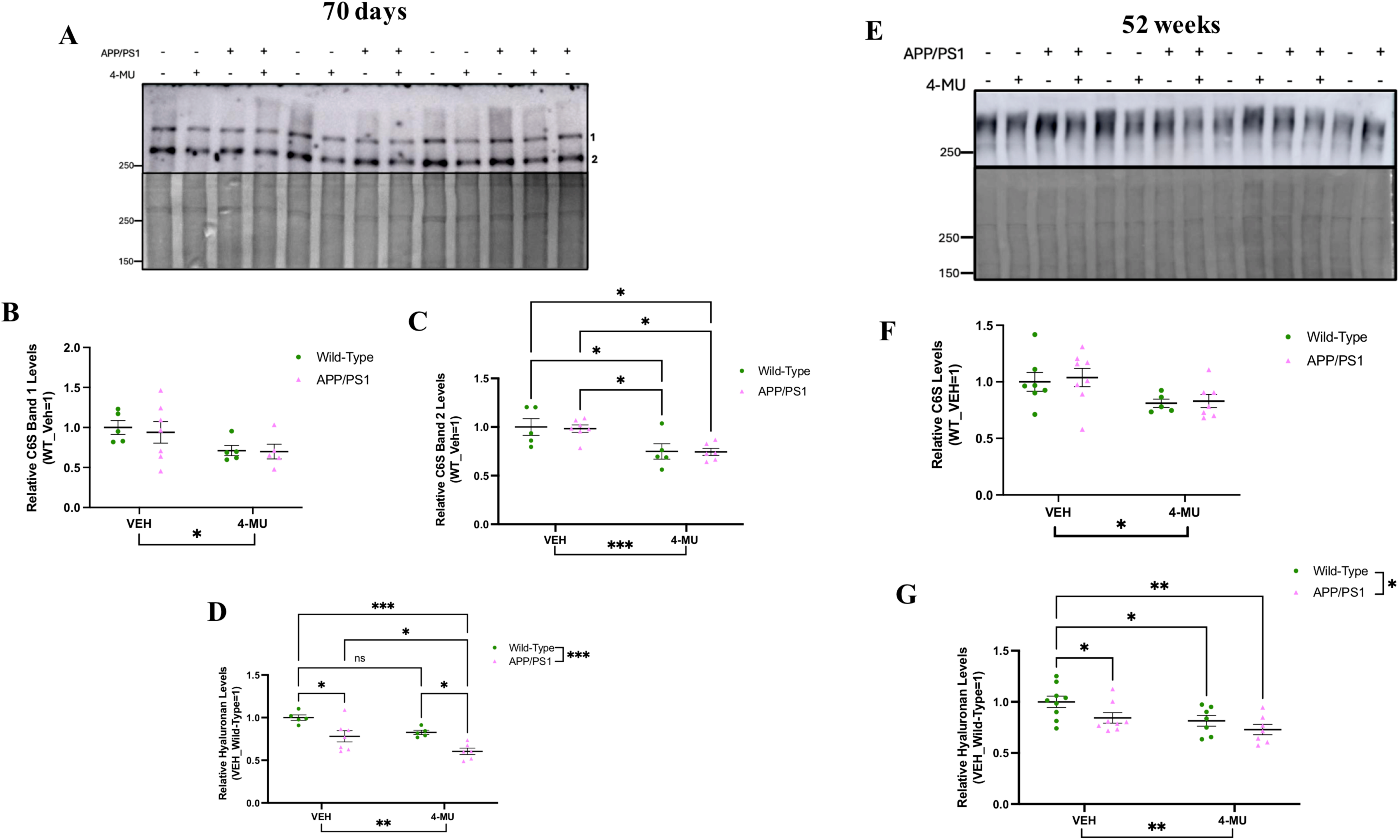
4-MU treatment reduces hippocampal hyaluronan and chondroitin-6-sulfate levels *in vivo*. **(A-C)** Western blot and densitometry showing 70 days of 4-MU treatment reduces two high molecular weight C6S bands. One outlier removed in APP/PS1 (4-MU) group in **B**. **(D)** ELISA showing 70 days of 4-MU treatment reduces hippocampal hyaluronic acid levels and 5.5-month-old APP/PS1 mice have less hyaluronan levels than WT littermates. 2-way ANOVA with Holm-Šidák’s multiple comparisons test. n=6-8 per group, mean ± SEM,***p<0.001, **p<0.01 *p<0.05. ROUT’s testing (Q=1%) used for outlier removal in **B-D**. **(E&F)** Western blot and densitometry showing 52 weeks of 4-MU treatment reduces a high molecular weight C6S. **(G)** 52 weeks of 4-MU treatment reduces hippocampal hyaluronan levels and 15-month-old APP/PS1 mice show less hyaluronan levels than WT littermates. 2-way ANOVA with Holm-Šidák’s multiple comparisons test. n=6-10 per group, mean ± SEM, **p<0.01, *p<0.05 in **F&G**.

Western blot analysis was used to assess PNN components in hippocampal lysates following 70 days and 52 weeks of 4-MU treatment. No significant changes were observed in hippocampal levels of aggrecan, neurocan, or versican at either time point (data not shown). In contrast, full-length brevican and its proteolytic fragments were increased following both treatment durations **(Figure 4)**. After 70 days of 4-MU treatment, levels of full-length brevican [F(1,19)= 5.383, p= 0.0316], 80-kDa C-terminal brevican [F(1,19)= 5.742, p= 0.027], and 55-kDa N-terminal brevican [F(1,19)= 6.453, p= 0.02] were significantly elevated relative to vehicle controls. Similar increases were observed after 52 weeks of treatment, with full-length brevican [F(1,23)= 11.26, p= 0.0027], 80-kDa brevican-C [F(1,23)= 12.79, p= 0.0016], and 55-kDa brevican-N [F(1,24)= 15.62, p= 0.0006] all significantly upregulated. HAPLN2, a link protein that anchors brevican and versican to HA in specialized perinodal matrices ^27^, was also assessed. Consistent with previous observations of elevated HAPLN2 in human *APOE4* targeted replacement mice and its localization in PNNs ^28^, 52 weeks of 4-MU treatment resulted in a significant reduction in prefrontal cortical HAPLN2 levels [F(1,24) = 4.722, p = 0.0399] and a trend toward decreased hippocampal HAPLN2 levels [F(1,24)= 2.988, p= 0.0967] **(Figure 5)**.

**Figure 4:**
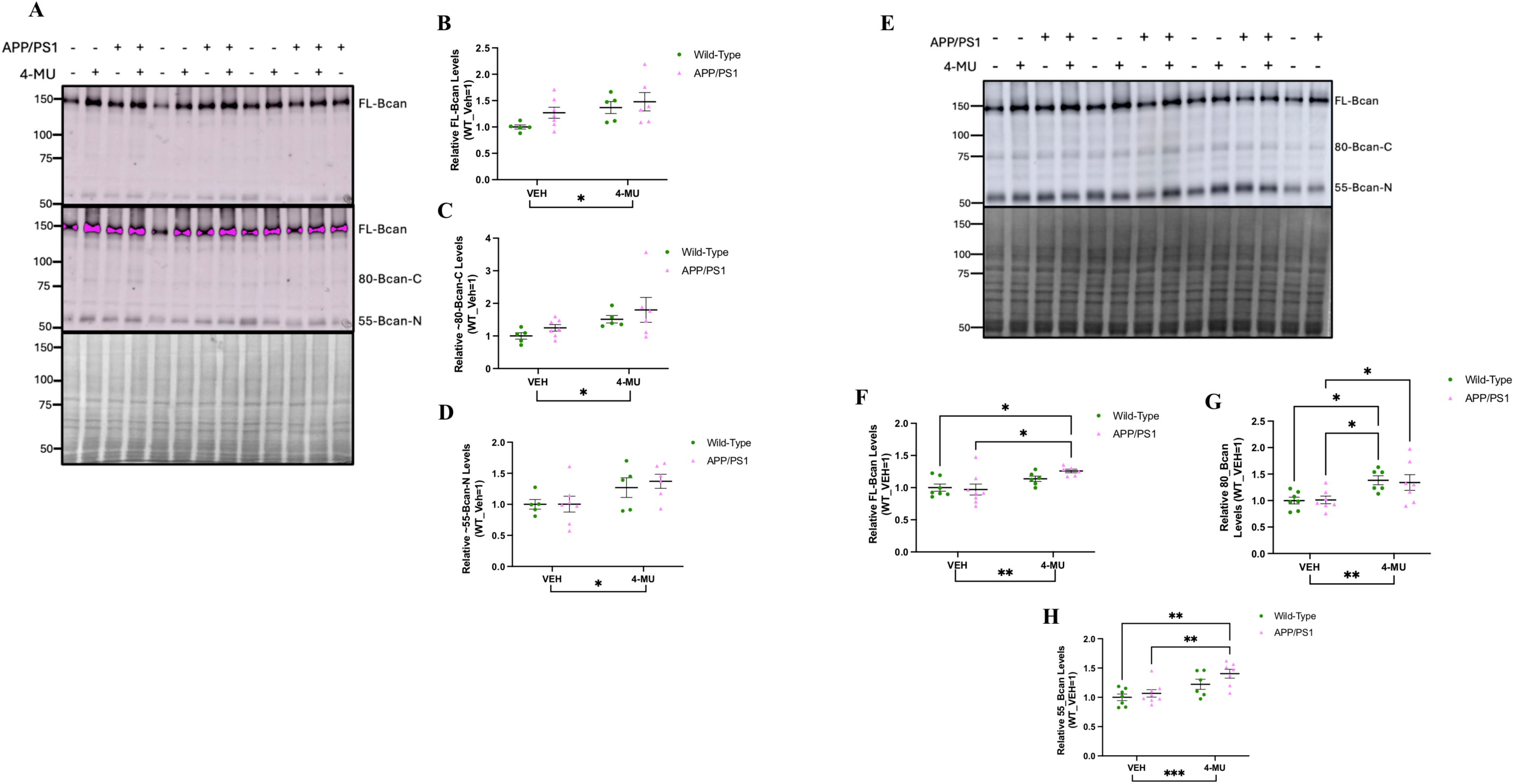
Short and long-term 4-MU treatment increases hippocampal brevican levels. **(A-D)** Western blot and densitometry showing that 70 days of 4-MU treatment increases brevican protein levels. Mean ± SEM, n=5-7 per group, *p<0.05. 2-way ANOVA with Holm-Šidák’s multiple comparisons test in **C&D**. **(E-H)** Western blot and densitometry showing that 52 weeks of 4-MU treatment increases brevican protein levels. One outlier removed in APP/PS1 (4-MU) in **F** and one outlier removed in APP/PS1 (VEH) in **G**. 2-way ANOVA with Holm-Šidák’s multiple comparisons test. n=6-8 per group, mean ± SEM, ***p<0.001, **p<0.01, *p<0.05, ROUT’s testing (Q=1%) used for outlier removal in **F-H**.

**Figure 5:**
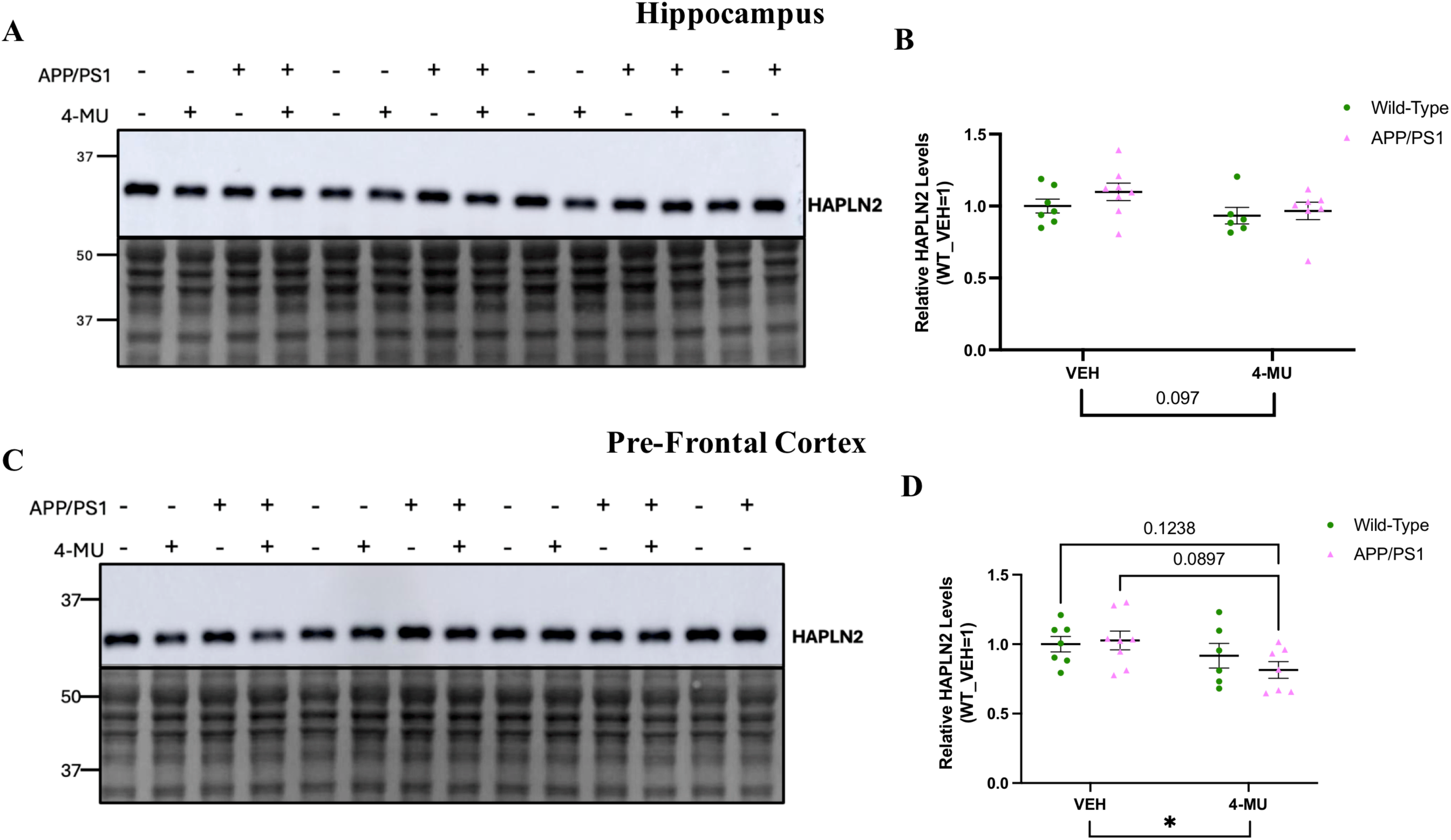
52 weeks of 4-MU treatment reduces pre-frontal cortical HAPLN2 levels and trends towards a reduction in the hippocampus. **(A&B)** Western blot and densitometry showing that 4-MU treatment trends towards a reduction in hippocampal HAPLN2 protein levels. **(C&D)**. Western blot and densitometry showing that 4-MU significantly reduces cortical HAPLN2 protein levels. 2-way ANOVA with Holm-Šidák’s multiple comparisons test. n=6-8 per group, mean ± SEM, *p<0.05.

To determine the effect of chronic 4-MU treatment on PNNs, we utilized IHC to measure the fluorescent intensity of hippocampal PNNs with two markers (WFA and aggrecan). WFA recognizes the non-sulfated chondroitin units on CSPGs, while the aggrecan antibody binds the core protein. We chose aggrecan over the other lecticans, because it has been shown to be indispensable for PNN formation and levels reflect PNN maturity ^29^. WFA is a conventional PNN marker; however, recent work shows it has little affinity for sulfated chondroitin isomers and high affinity for the non-sulfated chondroitin isomer ^30^. Furthermore, it was recently shown that total CS levels are elevated in AD postmortem brains ^31^, which emphasizes the need to use additional PNN markers instead of solely WFA. Previously it has been shown that 6 months of 4-MU treatment in WT mice reduces PNN fluorescent intensity ^18^. We found a significant decrease in PNN intensity across the 4 hippocampal subregions with both 70 days and 52 weeks of 4-MU treatment **(Figure 6)**. 70 days of 4-MU treatment resulted in a significant decrease in dentate gyrus aggrecan [F(1,18)= 9.274, p= 0.007] and PV [F(1,18)= 9.911, p= 0.0056] expression. 70 days of 4-MU treatment significantly decreased CA1 WFA [F(1,18)= 21.62, p= 0.0002], aggrecan [F(1,18)= 13.84, p= 0.0016], and PV [F(1,18)= 5.057, p= 0.0373] expression. 70 days of 4-MU treatment significantly decreased CA2 WFA [F(1,18)= 4.649, p= 0.0449] and aggrecan [F(1,18)= 8.280, p= 0.01] expression. 70 days of 4-MU treatment significantly decreased CA3 WFA [F(1,18)= 10.9, p= 0.004] and aggrecan [F(1,18)= 5.279, p= 0.0338] expression. 52 weeks of 4-MU treatment significantly reduced dentate gyrus WFA [F(1,20)= 5.576, p= 0.0285] and PV [F(1,20)= 4.94, p= 0.0379] expression. 52 weeks of 4-MU treatment significantly reduced CA1 WFA [F(1,20)= 10.12, p= 0.0047] and aggrecan [F(1,20)= 8.508, p= 0.0085] expression. 52 weeks of 4-MU treatment significantly reduced CA2 aggrecan expression [F(1,20)= 5.488, p= 0.0296]. 52 weeks of 4-MU treatment significantly reduced CA3 WFA [F(1,20)= 9.464, p= 0.006] and aggrecan [F(1,20)= 11.28, p= 0.0031] expression. No difference in PNN fluorescent intensity across the four hippocampal subregions was found between APP/PS1 mice and their WT littermates (p-values <0.05).

**Figure 6:**
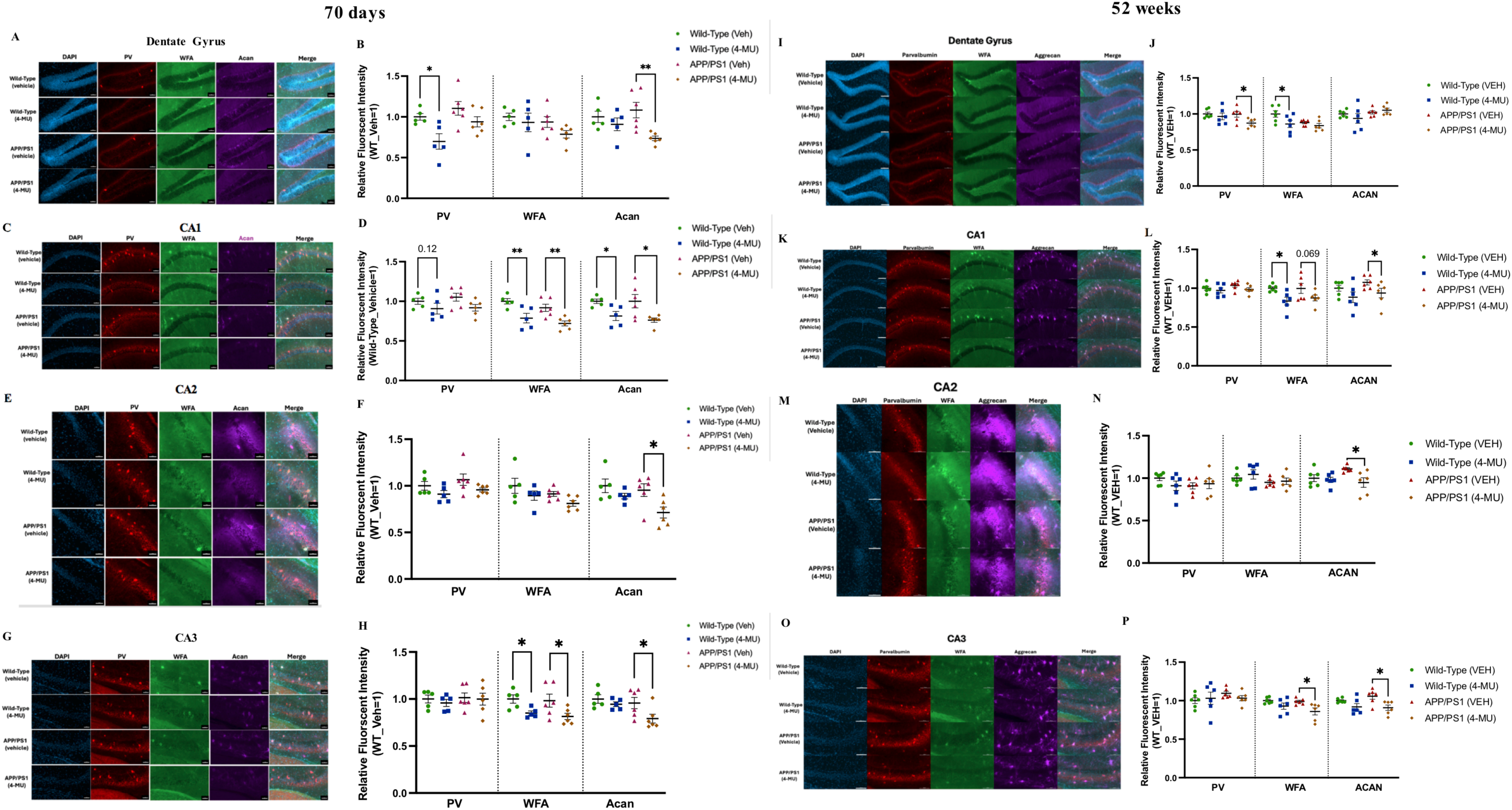
70 days and 52 weeks of 4-MU treatment reduces hippocampal PNN fluorescent intensity as measured with two markers, WFA and aggrecan. **(A&B)** 70 days of 4-MU treatment reduces dentate gyrus aggrecan and PV intensity. **(C&D)** 70 days of 4-MU treatment reduces CA1 WFA, aggrecan, and PV fluorescent intensity. **(E&F)** 70 days of 4-MU treatment reduces CA2 aggrecan intensity. **(G&H)** 70 days of 4-MU treatment reduces CA3 WFA and aggrecan fluorescent intensity. **(A-H)** Mean ± SEM, n=5-7 per group, **p<0.01, *p<0.05. 2-way ANOVA with Holm-Šidák’s multiple comparisons test. Scale bar= 50 µm. **(I&J)** 52 weeks of 4-MU treatment reduces dentate gyrus WFA and PV fluorescent intensity. **(K&L)** 52 weeks of 4-MU treatment reduces CA1 WFA and aggrecan fluorescent intensity. **(M&N)** 52 weeks of 4-MU treatment reduces CA2 aggrecan fluorescent intensity. **(O&P)** 52 weeks of 4-MU treatment reduces WFA and aggrecan fluorescent intensity. **(I-P)** Mean ± SEM, n=6 mice per group, **p<0.01, *p<0.05. 2-way ANOVA with Holm-Šidák’s multiple comparisons test. Scale bar= 100 µm.

### 4-MU attenuates Aβ pathology

APP/PS1 mice do not begin depositing insoluble amyloid plaque until ∼4 months of age, but soluble Aβ species are present at appreciable levels ^32,33^. Furthermore, mounting evidence implicates the soluble Aβ oligomers in driving neuroinflammation and subsequent neuronal loss ^34,35^. We measured soluble Aβ species (Aβ_1-40_ and Aβ_1-42_ ) in mouse brain tissue pre-frontal cortical and hippocampal lysates with ELISAs **(Figure 7)**. We found no effect of 70 days of 4-MU treatment on cortical [t(11)= 1.695, p= 0.1182] and hippocampal [(t11)= 0.3353, p= 0.7437] soluble Aβ_1-40_ levels and cortical [t(11)= 0.148, p= 0.885] and hippocampal Aβ_1-42_ levels [t(11)= 1.258, p= 0.2345]. 52 weeks of 4-MU treatment did not affect cortical [t(13)= 0.4899, p=0.6324] and hippocampal [t(13)= 0.1601, p= 0.8753] soluble Aβ_1-40_ levels but trended towards a reduction in cortical soluble Aβ_1-42_levels [t(13)= 1.915, p=0.0778]. We noted a significant reduction in the parenchymal Aβ_1-42_/Aβ_1-40_ ratio with both 70 days [F(1,22)= 10.68, p= 0.0035] and 52 weeks [F(1,26)= 5.675, p= 0.0248] of 4-MU treatment. Furthermore, the pre-frontal cortex demonstrated a greater Aβ_1-42_/Aβ_1-40_ ratio than the hippocampus in both the 5.5-month [F(1,22)= 9.008, p=0.0066] and 15-month [F(1,26)= 45.92, p< 0.0001] old APP/PS1 mice. We measured insoluble amyloid plaque levels in the cortex with IF using the 6E10 antibody which binds the N-terminus of humanized Aβ species **(Figure 8)**. Notably, we found that 52 weeks of 4-MU treatment reduced cortical insoluble amyloid plaque levels [t(11)= 3.103, p= 0.01]. Taken together, this data lends support for HA inhibition in the treatment of AD.

**Figure 7:**
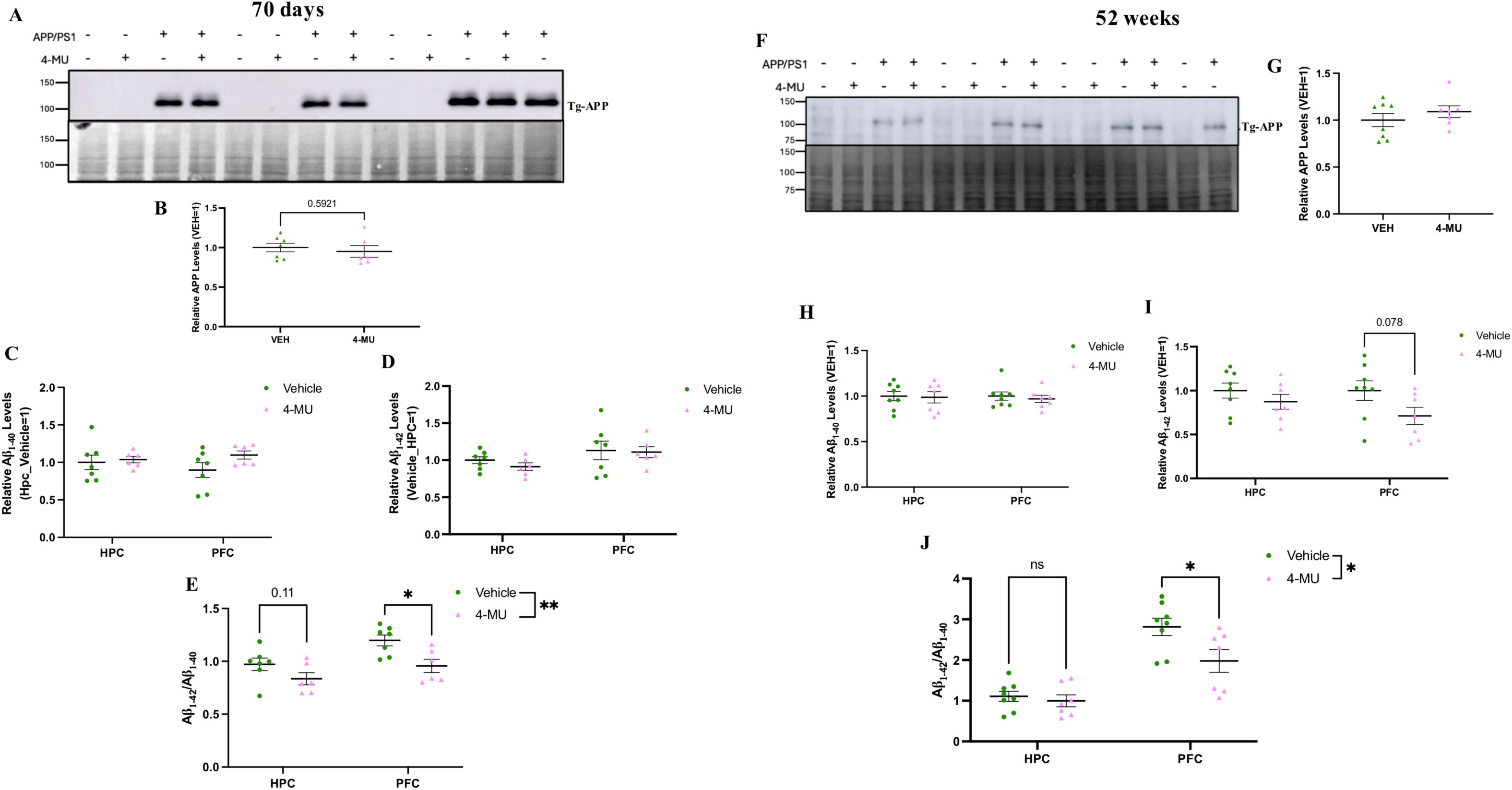
70 days and 52 weeks of 4-MU treatment reduces the parenchymal Aβ_1-42_/Aβ_1-40_ ratio. **(A&B)** Western blot and densitometry showing that 70 days of 4-MU treatment does not affect hippocampal transgenic APP levels. **(C&D)** 70 days of 4-MU treatment does not affect cortical and hippocampal soluble levels of Aβ_1-40_ and Aβ_1-42_. **(E)** 70 days of 4-MU treatment reduces the brain parenchymal Aβ_1-42_/Aβ_1-40_ratio. n=6-8 per group, mean ± SEM, student’s unpaired two tailed T test in **A-D** and two-way ANOVA with Holm-Šidák’s multiple comparisons test in **E**, **p<0.01, *p<0.05. **(F&G)** Western blot and densitometry showing 52 weeks of 4-MU treatment does not affect hippocampal transgenic APP levels. **(H&I) )** 52 weeks of 4-MU treatment does not affect hippocampal soluble levels of Aβ_1-40_and Aβ_1-42_ but trends towards a reduction in pre-frontal cortical Aβ_1-42_ levels. **(J)** 52 weeks of 4-MU treatment reduces the brain parenchymal Aβ_1-42_/Aβ_1-40_ ratio. n=7-8 per group, mean ± SEM, student’s unpaired two tailed T test in **F-I** and two-way ANOVA with Holm-Šidák’s multiple comparisons test in **J**, *p<0.05.

**Figure 8:**
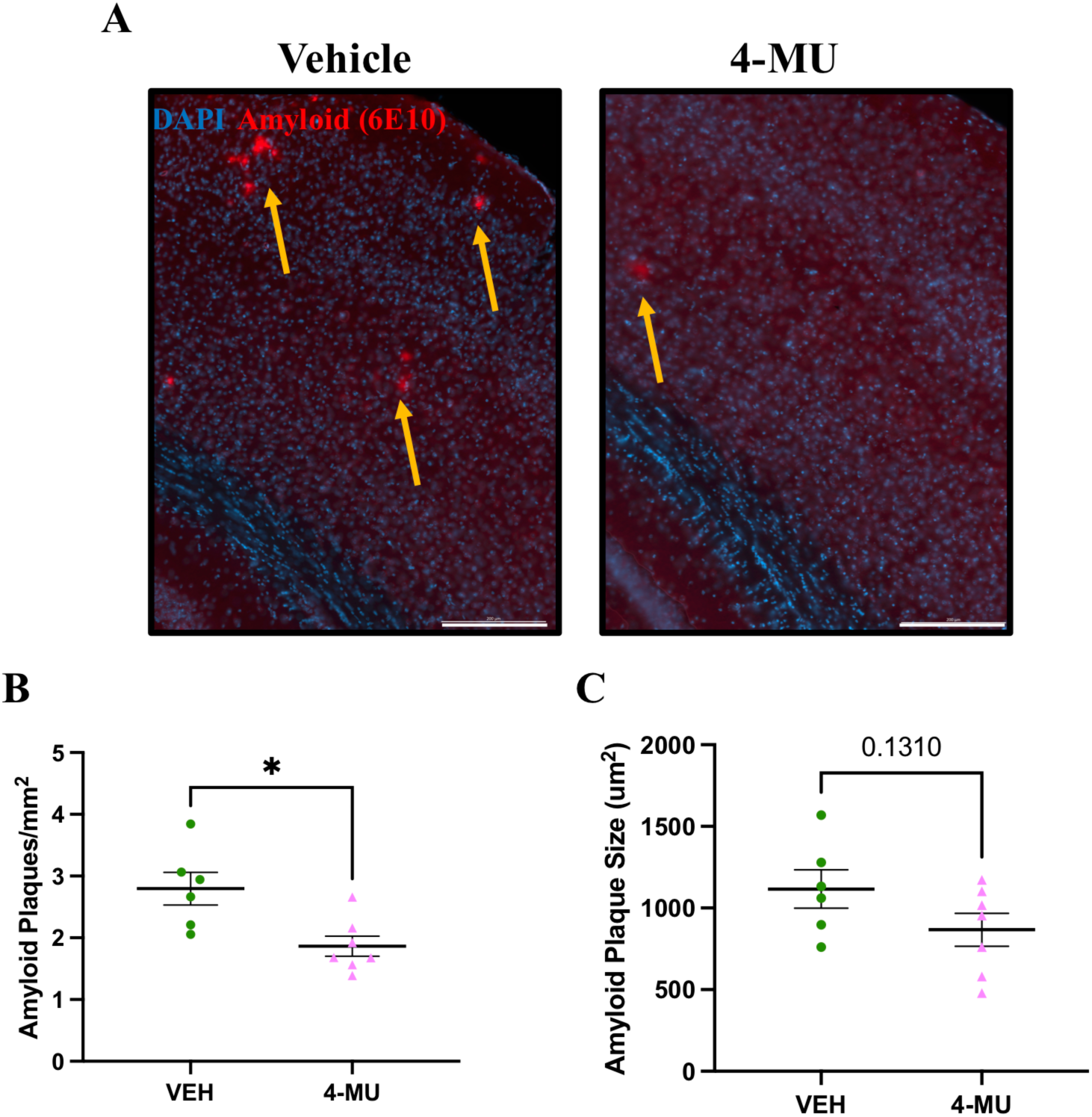
52 weeks of 4-MU treatment reduces cortical insoluble amyloid plaque levels. **(A)** Representative images of amyloid plaque in the cortex of vehicle and 4-MU treated mice. 6E10 antibody was used. **(B)** Cortical amyloid plaque levels and **(C)** cortical amyloid plaque size. Yellow arrows indicate amyloid plaques Student’s unpaired two-tailed T test, mean ± SEM, *p<0.05, n=6-7 per group. Scale bar = 200 µm.

## Discussion

Herein, we investigated the effects of 4-MU on Aβ pathology, ECM/PNN dynamics, and cognition in the APP/PS1 mouse model of AD. Prior work has highlighted a complex relationship between Aβ pathology and PNN components. Brevican has been shown to bind Aβ species *in vitro* ^11^ and to co-localize with amyloid plaques *in vivo* ^39^. In addition, tenascin-C is enriched within dense-core amyloid plaques in postmortem AD brain tissue ^40^. Conversely, overall PNN integrity is reduced in some amyloidogenic models, including 5xFAD mice ^41,42^. Importantly, however, several studies have demonstrated that enzymatic degradation of PNNs in both amyloid- ^11–13^ and tau- ^43^ based mouse models attenuates AD-related neuropathology. Herein, we assessed the therapeutic potential of chronic pharmacological inhibition of PNN formation using 4-MU in an amyloid mouse model. Notably, both short- and long-term 4-MU treatment resulted in a reduction of Aβ pathology. 70 days of 4-MU administration significantly reduced the cortical soluble Aβ_1-42_/Aβ_1-40_ ratio. 52 weeks of treatment led to reductions in both the soluble Aβ_1-42_/Aβ_1-40_ ratio and insoluble amyloid plaque levels. The reduction in the parenchymal Aβ_1-42_/Aβ_1-40_ ratio is noteworthy, as this ratio increases with disease progression in individuals with familial AD ^44,45^. Recent evidence also indicates that elevated Aβ_1-42_/Aβ_1-40_ ratios within soluble protofibrils strongly predict reduced CSF Aβ_1-42_/Aβ_1-40_ ratios, as well as increased CSF neurofilament light and tau levels ^46^.

A caveat of our study is that we did not examine potential effects of 4-MU on heparan sulfate proteoglycan (HSPG) components of brain ECM. Because of their role in memory as well as neuronal oscillations that are impaired in AD ^47^, our initial focus was on CSPGs. Of interest, however, 4-MU also has the potential to reduce the protein expression of select HSPGs. This is because UDP-glucuronic acid is also a substrate for heparan sulfate synthesis ^48,49^. Importantly, HSPGs have also been implicated in amyloid aggregation and may impact glymphatic clearance of toxic proteins ^50–53^.

With respect to potential therapeutic use, chronic 4-MU treatment was well tolerated and appeared safe in both APP/PS1 mice and controls. Furthermore, a recently published study showed that chronic administration of 5% 4-MU in diet increased the average lifespan of WT mice ^23^. 4-MU–treated mice exhibited an approximate 16% reduction in body weight; however, treated animals continued to gain weight with age. Prior studies have also reported 4-MU–induced reductions in body weight ^20,54^. 4-MU has been shown to lower blood glucose levels and enhance insulin sensitivity in mouse models of metabolic disease ^22,23^. Since type 2 diabetes and obesity represent two modifiable risk factors for AD, this metabolic phenotype may be advantageous ^55^.

Despite the abundance of HA in the brain and PNNs, HA is highly concentrated in cartilage-rich tissues. Previous work has demonstrated that six months of 4-MU treatment in mice does not impair motor function, as assessed by a lower speed rotarod assay and grip strength ^18^. In contrast, we observed that 70 days but not 52 weeks of 4-MU treatment was associated with reduced performance on the rotarod task. These findings suggest that short-term 4-MU exposure may transiently impair motor coordination or endurance, with recovery occurring over longer post developmental treatment durations.

Spatial memory is impaired in individuals with AD and in transgenic mouse models of AD ^56,57^. Accordingly, we evaluated the effects of 52 weeks of 4-MU treatment on spatial learning and memory using the Barnes maze in control and APP/PS1 mice. We also incorporated a reversal-learning paradigm to assess cognitive flexibility, a domain known to be constrained by PNN–mediated stabilization of circuits. Across genotypes, 4-MU-treated mice exhibited reduced average escape latencies during the acquisition phase of the standard Barnes maze compared with vehicle-treated controls, indicating enhanced spatial learning. Notably, this effect was also observed in healthy WT mice, suggesting that long-term 4-MU treatment can improve learning even in the absence of amyloid pathology. This finding is consistent with prior reports demonstrating age-related impairments in Barnes maze performance ^58^ and evidence that 4-MU enhances object recognition memory in adult WT mice ^18^. 4-MU treatment did not significantly affect performance on the probe trial, a measure of spatial recall and long-term memory retention, indicating that the benefits of 4-MU may preferentially impact acquisition rather than memory consolidation or retrieval. Importantly, 4-MU–treated mice, regardless of genotype, demonstrated improved performance during the acquisition phase of the reversal-learning component of the Barnes maze, consistent with increased cognitive flexibility.

With respect to ECM related endpoints, 4-MU treatment resulted in an expected and significant reduction of hippocampal HA levels following both short and long-term administration. Hippocampal HA levels were reduced in both 5.5- and 15-month-old APP/PS1 mice relative to age-matched controls, suggesting that amyloid pathology itself is associated with alterations in HA homeostasis. HA is a major constituent of the diffuse ECM, where it contributes to the regulation of neuronal excitability ^59^ and glial inflammatory responses ^60^. Genetic depletion of HA has been shown to induce spontaneous epileptiform activity and reduce ECS volume in the brain ^61^. Importantly, no treatment-associated mortality was observed in the present study, suggesting that any proconvulsant effects of 4-MU, if present, were limited under the experimental conditions employed. Future investigations with longitudinal electroencephalographic (EEG) monitoring will be required to better examine this issue.

Both 70 days and 52 weeks of 4-MU treatment also resulted in a decrease in hippocampal C6S levels in APP/PS1 and WT mice; however, no effect of 4-MU treatment was observed on C4S levels. C6S levels are relatively increased during early development and levels are reduced with aging and synaptic maturation ^62^. C6S generally favors neuroplasticity and overexpression of C6S increases PNN proteolytic degradation ^63^ and improves object recognition memory in aged mice ^64^. Interestingly, however, temporal cortex and hippocampal C6S levels are elevated in AD humans and they positively correlate with Braak stage pathology and p-tau 396 levels ^31^. Furthermore, C6S levels were shown to be the most predictive CS isomer of dementia burden in humans ^65^.

We also observed a reduction in pre-frontal cortical HAPLN2 levels with 52 weeks of 4-MU treatment. HAPLN2 forms insoluble aggregates in the brain parenchyma which may increase with age ^66^ and expression of APOE4 ^28^. It may be of interest to determine if insoluble HAPLN2 deposits associate with amyloid plaques.

We also found an increase in hippocampal brevican levels with both 70 days and 52 weeks of 4-MU treatment. Our findings show that 4-MU administration led to elevated levels of full length brevican. Dubisova and colleagues previously reported a trend toward increased *brevican* mRNA expression in WT mice following prolonged 4-MU treatment ^18^, and the concurrent elevation of both intact and cleaved brevican proteins observed here supports the possibility that 4-MU enhances brevican synthesis at the transcriptional level. Further work will be necessary to define the molecular pathways underlying 4-MU–mediated regulation of CSPG gene expression.

4-MU treatment also reduced hippocampal PNN fluorescent intensity across hippocampal subregions. The reductions detected in both WFA- and aggrecan-positive PNNs indicate that 4-MU induces remodeling of hippocampal PNN architecture. Intact PNNs are required to sustain the fast-spiking properties of PV neurons, and enzymatic or structural disruption of these nets has been shown to reduce PV neuronal activity ^47^. Attenuation of PNNs by 4-MU may secondarily alter PV interneuron function, with potential consequences for hippocampal network dynamics.

Interestingly, we found that 4-MU administration to primary glia reduces basal and TGF-β1-stimulated CS levels. Previously it was shown that chronic 4-MU treatment in WT mice reduces PNNs labeled with a CS marker ^18^, and our data further supports 4-MU as a direct CS inhibitor. 4-MU may thus have the ability to reduce PNNs independent of HA inhibition.

In summary, we show that chronic 4-MU treatment in an amyloid mouse model improves learning and reduces Aβ pathology. To our knowledge, this is the first study to assess long-term 4-MU administration in a neurodegenerative disease model, and it supports the potential for HA inhibition to reduce Aβ pathology. Future studies should examine 4-MU effects in other AD models and its effects on ECM remodeling at other sites including the blood brain barrier which may be damaged following anti-Aβ immunotherapy ^67^.

## Acknowledgements

The authors would like to acknowledge support from the National Institutes of Health (AG077002; KC).

## Disclosures/Conflicts of Interest

The authors have nothing to disclose and declare no conflicts of interest.

## Supplemental Figures

**Supplemental Figure 1:** 70 days and 52 weeks of 4-MU treatment reduce body weight in both male and female mice and do not affect food consumption. No difference in body weight was found between APP/PS1 and WT mice, so they are grouped in this analysis. **(A&B)** 70 days of 4-MU treatment reduces body weight in male and female mice. **(C&D)** 70 days of 4-MU treatment does not affect daily food consumption. n=5-6 per group in **A&B** and n=2 cages per group in **C&D**, mean ± SEM, 2-way repeated measures ANOVA, **p<0.01, *p<0.05. **(E&F)** 52 weeks of 4-MU treatment reduces body weight in both male and female mice. **(G&H)** 52 weeks of 4-MU treatment does not affect food consumption in female mice but trends towards a reduced consumption in male mice. n=6-9 per group in **E&F** and n=2 cages per group in **G&H**, mean ± SEM, 2-way repeated measures ANOVA.

**Supplemental Figure 2:** 70 days but not 52 weeks of 4-MU treatment reduces performance on the rotarod behavioral assay. **(A)** Graphical schematic of rotarod apparatus. Created in BioRender. **(B)** 70 days of 4-MU treatment reduces latency to fall on the rotarod for both APP/PS1 mice and WT littermates. 2-way ANOVA with Holm-Šidák’s multiple comparisons test. n=5-7 per group, mean ± SEM, **p<0.01, *p<0.05. **(C)** 52 weeks of 4-MU treatment does not affect performance on rotarod assay. 15-month-old APP/PS1 mice trended towards a reduction in latency to fall on the rotarod. 2-way ANOVA with Holm-Šidák’s multiple comparisons test. n=7-9 per group Mean ± SEM.

**Supplemental Figure 3:** Previously it has been shown that 4-MU blunts astrocyte activation and subsequent prostaglandin and cytokine release in response to LPS *in vitro* ^36^ and *in vivo* ^37^. Because 4-MU inhibits UDP-glucuronic acid formation, a substrate also required for CS synthesis, and has also been reported to reduce CS-labeled PNNs ^18^, we tested whether 4-MU affects CS levels in primary glial cultures. 24 hours of 4-MU exposure in primary glia reduces HA and chondroitin-4-sulfate (C4S) levels. **(A)** ELISA showing 24 hours of 4-MU exposure reduces the levels of HA secreted into the media by astrocytes. **(B-D)** Western blot and densitometry showing 24 hours of 4-MU treatment decreases high molecular weight C4S levels in the glial cell lysates at basal conditions and with TGF-β1 stimulation. Mean ± SEM, n=3 technical replicates per group, n=2 independent experiments performed. A representative experiment is shown. ***p<0.001, **p<0.01, *p<0.05. 2-way ANOVA with Holm-Šidák’s multiple comparisons test. 4-MU dose used in these experiments was 500 µM and was administered 1 hour prior to TGF-β1 (10 ng/mL) treatment.

## Notes

### Competing Interest Statement

The authors have declared no competing interest.

## References

1. Tiwari S, Atluri V, Kaushik A, Yndart A, Nair M. Alzheimer’s disease: pathogenesis, diagnostics, and therapeutics. Int J Nanomedicine. 2019;14:5541–5554. doi:10.2147/IJN.S200490

2. Zheng Q, Wang X. Alzheimer’s disease: insights into pathology, molecular mechanisms, and therapy. Protein Cell. 2025;16(2):83–120. doi:10.1093/procel/pwae026

3. Xu H, Garcia-Ptacek S, Jönsson L, Wimo A, Nordström P, Eriksdotter M. Long-term Effects of Cholinesterase Inhibitors on Cognitive Decline and Mortality. Neurology. 2021;96(17):e2220–e2230. doi:10.1212/WNL.0000000000011832

4. Bateman RJ, Xiong C, Benzinger TLS, et al. Clinical and Biomarker Changes in Dominantly Inherited Alzheimer’s Disease. N Engl J Med. 2012;367(9):795–804. doi:10.1056/NEJMoa1202753

5. Ulbrich P, Khoshneviszadeh M, Jandke S, Schreiber S, Dityatev A. Interplay between perivascular and perineuronal extracellular matrix remodelling in neurological and psychiatric diseases. Eur J Neurosci. 2021;53(12):3811–3830. doi:10.1111/ejn.14887

6. Lupori L, Totaro V, Cornuti S, et al. A comprehensive atlas of perineuronal net distribution and colocalization with parvalbumin in the adult mouse brain. Cell Rep. 2023;42(7):112788. doi:10.1016/j.celrep.2023.112788

7. Hensch TK. Critical period plasticity in local cortical circuits. Nat Rev Neurosci. 2005;6(11):877–888. doi:10.1038/nrn1787

8. Pizzorusso T, Medini P, Berardi N, Chierzi S, Fawcett JW, Maffei L. Reactivation of ocular dominance plasticity in the adult visual cortex. Science. 2002;298(5596):1248–1251. doi:10.1126/science.1072699

9. Carulli D, Pizzorusso T, Kwok JCF, et al. Animals lacking link protein have attenuated perineuronal nets and persistent plasticity. Brain J Neurol. 2010;133(Pt 8):2331–2347. doi:10.1093/brain/awq145

10. Gogolla N, Caroni P, Lüthi A, Herry C. Perineuronal nets protect fear memories from erasure. Science. 2009;325(5945):1258–1261. doi:10.1126/science.1174146

11. Howell MD, Bailey LA, Cozart MA, Gannon BM, Gottschall PE. Hippocampal administration of chondroitinase ABC increases plaque-adjacent synaptic marker and diminishes amyloid burden in aged APPswe/PS1dE9 mice. Acta Neuropathol Commun. 2015;3(1):54. doi:10.1186/s40478-015-0233-z

12. Végh MJ, Heldring CM, Kamphuis W, et al. Reducing hippocampal extracellular matrix reverses early memory deficits in a mouse model of Alzheimer’s disease. Acta Neuropathol Commun. 2014;2:76. doi:10.1186/s40478-014-0076-z

13. Yang Q, Yan C, Sun Y, et al. Extracellular Matrix Remodeling Alleviates Memory Deficits in Alzheimer’s Disease by Enhancing the Astrocytic Autophagy-Lysosome Pathway. Adv Sci Weinh Baden-Wurtt Ger. 2024;11(31):e2400480. doi:10.1002/advs.202400480

14. Kultti A, Pasonen-Seppänen S, Jauhiainen M, et al. 4-Methylumbelliferone inhibits hyaluronan synthesis by depletion of cellular UDP-glucuronic acid and downregulation of hyaluronan synthase 2 and 3. Exp Cell Res. 2009;315(11):1914–1923. doi:10.1016/j.yexcr.2009.03.002

15. Kakizaki I, Kojima K, Takagaki K, et al. A Novel Mechanism for the Inhibition of Hyaluronan Biosynthesis by 4-Methylumbelliferone*. J Biol Chem. 2004;279(32):33281–33289. doi:10.1074/jbc.M405918200

16. Galgoczi E, Jeney F, Katko M, et al. Characteristics of Hyaluronan Synthesis Inhibition by 4-Methylumbelliferone in Orbital Fibroblasts. Invest Ophthalmol Vis Sci. 2020;61(2):27. doi:10.1167/iovs.61.2.27

17. Sukowati CHC, Anfuso B, Fiore E, et al. Hyaluronic acid inhibition by 4-methylumbelliferone reduces the expression of cancer stem cells markers during hepatocarcinogenesis. Sci Rep. 2019;9(1):4026. doi:10.1038/s41598-019-40436-6

18. Dubisova J, Burianova JS, Svobodova L, et al. Oral treatment of 4-methylumbelliferone reduced perineuronal nets and improved recognition memory in mice. Brain Res Bull. 2022;181:144–156. doi:10.1016/j.brainresbull.2022.01.011

19. Štepánková K, Chudíčková M, Šimková Z, et al. Low oral dose of 4-methylumbelliferone reduces glial scar but is insufficient to induce functional recovery after spinal cord injury. Sci Rep. 2023;13(1):19183. doi:10.1038/s41598-023-46539-5

20. Grandoch M, Flögel U, Virtue S, et al. 4-Methylumbelliferone improves the thermogenic capacity of brown adipose tissue. Nat Metab. 2019;1(5):546–559. doi:10.1038/s42255-019-0055-6

21. Nagy N, Kaber G, Sunkari VG, et al. Inhibition of hyaluronan synthesis prevents β-cell loss in obesity-associated type 2 diabetes. Matrix Biol. 2023;123:34–47. doi:10.1016/j.matbio.2023.09.003

22. Zhang W, Yang S, Yu X, et al. Beneficial Actions of 4-Methylumbelliferone in Type 1 Diabetes by Promoting β Cell Renewal and Inhibiting Dedifferentiation. Biomedicines. 2024;12(12):2790. doi:10.3390/biomedicines12122790

23. Nagy N, Czepiel KS, Kaber G, et al. Hymecromone Promotes Longevity and Insulin Sensitivity in Mice. Cells. 2024;13(20):1727. doi:10.3390/cells13201727

24. Syková E, Voříšek I, Starčuk Z, et al. Disruption of extracellular matrix and perineuronal nets modulates extracellular space volume and geometry. J Neurosci. Published online January 2, 2025. doi:10.1523/JNEUROSCI.0517-24.2024

25. Coughlan G, Laczó J, Hort J, Minihane AM, Hornberger M. Spatial navigation deficits — overlooked cognitive marker for preclinical Alzheimer disease? Nat Rev Neurol. 2018;14(8):496–506. doi:10.1038/s41582-018-0031-x

26. Zhu H, Yan H, Tang N, et al. Impairments of spatial memory in an Alzheimer’s disease model via degeneration of hippocampal cholinergic synapses. Nat Commun. 2017;8(1):1676. doi:10.1038/s41467-017-01943-0

27. Oohashi T, Hirakawa S, Bekku Y, et al. Bral1, a brain-specific link protein, colocalizing with the versican V2 isoform at the nodes of Ranvier in developing and adult mouse central nervous systems. Mol Cell Neurosci. 2002;19(1):43–57. doi:10.1006/mcne.2001.1061

28. Deasy S, Amontree M, Colon Z, et al. Increased levels of HAPLN2, which anchors dense extracellular matrix, in the hippocampus of APOE4 targeted replacement mice. BioRxiv Prepr Serv Biol. Published online November 10, 2025:2025.11.09.687435. doi:10.1101/2025.11.09.687435

29. Rowlands D, Lensjø KK, Dinh T, et al. Aggrecan Directs Extracellular Matrix-Mediated Neuronal Plasticity. J Neurosci. 2018;38(47):10102–10113. doi:10.1523/JNEUROSCI.1122-18.2018

30. Scarlett JM, Hu SJ, Alonge KM. The “Loss” of Perineuronal Nets in Alzheimer’s Disease: Missing or Hiding in Plain Sight? Front Integr Neurosci. 2022;16:896400. doi:10.3389/fnint.2022.896400

31. Logsdon AF, Francis KL, Richardson NE, et al. Decoding perineuronal net glycan sulfation patterns in the Alzheimer’s disease brain. Alzheimers Dement J Alzheimers Assoc. 2022;18(5):942–954. doi:10.1002/alz.12451

32. Takeda S, Hashimoto T, Roe AD, Hori Y, Spires-Jones TL, Hyman BT. Brain interstitial oligomeric amyloid β increases with age and is resistant to clearance from brain in a mouse model of Alzheimer’s disease. FASEB J. 2013;27(8):3239–3248. doi:10.1096/fj.13-229666

33. Zhang L, Yang C, Li Y, et al. Dynamic Changes in the Levels of Amyloid-β42 Species in the Brain and Periphery of APP/PS1 Mice and Their Significance for Alzheimer’s Disease. Front Mol Neurosci. 2021;14. doi:10.3389/fnmol.2021.723317

34. Tolar M, Hey J, Power A, Abushakra S. Neurotoxic Soluble Amyloid Oligomers Drive Alzheimer’s Pathogenesis and Represent a Clinically Validated Target for Slowing Disease Progression. Int J Mol Sci. 2021;22(12):6355. doi:10.3390/ijms22126355

35. Zhang W, Hao J, Liu R, et al. Soluble Aβ levels correlate with cognitive deficits in the 12-month-old APPswe/PS1dE9 mouse model of Alzheimer’s disease. Behav Brain Res. 2011;222(2):342–350. doi:10.1016/j.bbr.2011.03.072

36. Chistyakov DV, Nikolskaya AI, Goriainov SV, Astakhova AA, Sergeeva MG. Inhibitor of Hyaluronic Acid Synthesis 4-Methylumbelliferone as an Anti-Inflammatory Modulator of LPS-Mediated Astrocyte Responses. Int J Mol Sci. 2020;21(21):8203. doi:10.3390/ijms21218203

37. Chistyakov DV, Nikolskaya AI, Gorbatenko VO, Goriainov SV, Silachev DN, Sergeeva MG. Inhibitor of hyaluronic acid synthesis 4-methylumbelliferone (4-MU) as a potential anti-inflammatory substance in acute neuroinflammation model in vivo. Inflammopharmacology. 2026;34(1):485–494. doi:10.1007/s10787-025-02063-8

38. Amontree M, Nelson M, Stefansson L, et al. Resveratrol differentially affects MMP-9 release from neurons and glia; implications for therapeutic efficacy. J Neurochem. Published online January 1, 2024. doi:10.1111/jnc.16031

39. Ajmo JM, Bailey LA, Howell MD, et al. Abnormal Post-Translational and Extracellular Processing of Brevican in Plaque-Bearing Mice Overexpressing APPsw. J Neurochem. 2010;113(3):784–795. doi:10.1111/j.1471-4159.2010.06647.x

40. Mi Z, Halfter W, Abrahamson EE, et al. Tenascin-C Is Associated with Cored Amyloid-β Plaques in Alzheimer Disease and Pathology Burdened Cognitively Normal Elderly. J Neuropathol Exp Neurol. 2016;75(9):868–876. doi:10.1093/jnen/nlw062

41. Barahona RA, Kwang NE, Kono-Soosaipillai AR, et al. Aggrecan protects against plaque accumulation and is essential for proper microglial responses to plaques. Cell Rep. 2025;44(8):116064. doi:10.1016/j.celrep.2025.116064

42. Crapser JD, Spangenberg EE, Barahona RA, Arreola MA, Hohsfield LA, Green KN. Microglia facilitate loss of perineuronal nets in the Alzheimer’s disease brain. EBioMedicine. 2020;58:102919. doi:10.1016/j.ebiom.2020.102919

43. Yang S, Cacquevel M, Saksida LM, et al. Perineuronal net digestion with chondroitinase restores memory in mice with tau pathology. Exp Neurol. 2015;265:48–58. doi:10.1016/j.expneurol.2014.11.013

44. Kumar-Singh S, Theuns J, Van Broeck B, et al. Mean age-of-onset of familial alzheimer disease caused by presenilin mutations correlates with both increased Abeta42 and decreased Abeta40. Hum Mutat. 2006;27(7):686–695. doi:10.1002/humu.20336

45. Scheuner D, Eckman C, Jensen M, et al. Secreted amyloid beta-protein similar to that in the senile plaques of Alzheimer’s disease is increased in vivo by the presenilin 1 and 2 and APP mutations linked to familial Alzheimer’s disease. Nat Med. 1996;2(8):864–870. doi:10.1038/nm0896-864

46. Andersson E, Lindblom N, Janelidze S, et al. Soluble cerebral Aβ protofibrils link Aβ plaque pathology to changes in CSF Aβ42/Aβ40 ratios, neurofilament light and tau in Alzheimer’s disease model mice. Nat Aging. 2025;5(3):366–375. doi:10.1038/s43587-025-00810-8

47. Lensjø KK, Lepperød ME, Dick G, Hafting T, Fyhn M. Removal of Perineuronal Nets Unlocks Juvenile Plasticity Through Network Mechanisms of Decreased Inhibition and Increased Gamma Activity. J Neurosci Off J Soc Neurosci. 2017;37(5):1269–1283. doi:10.1523/JNEUROSCI.2504-16.2016

48. Esko JD, Selleck SB. Order out of chaos: assembly of ligand binding sites in heparan sulfate. Annu Rev Biochem. 2002;71:435–471. doi:10.1146/annurev.biochem.71.110601.135458

49. Selva EM, Hong K, Baeg GH, et al. Dual role of the fringe connection gene in both heparan sulphate and fringe-dependent signalling events. Nat Cell Biol. 2001;3(9):809–815. doi:10.1038/ncb0901-809

50. Berzin TM, Zipser BD, Rafii MS, et al. Agrin and microvascular damage in Alzheimer’s disease. Neurobiol Aging. 2000;21(2):349–355. doi:10.1016/s0197-4580(00)00121-4

51. Holmes BB, DeVos SL, Kfoury N, et al. Heparan sulfate proteoglycans mediate internalization and propagation of specific proteopathic seeds. Proc Natl Acad Sci. 2013;110(33):E3138–E3147. doi:10.1073/pnas.1301440110

52. Verbeek MM, Otte-Höller I, Veerhuis R, Ruiter DJ, De Waal RM. Distribution of A beta-associated proteins in cerebrovascular amyloid of Alzheimer’s disease. Acta Neuropathol (Berl). 1998;96(6):628–636. doi:10.1007/s004010050944

53. Ye F, Li M, Liu M, et al. Co-Aggregation of Syndecan-3 with β-Amyloid Aggravates Neuroinflammation and Cognitive Impairment in 5×FAD Mice. Int J Mol Sci. 2025;26(12):5502. doi:10.3390/ijms26125502

54. Kuipers HF, Nagy N, Ruppert SM, et al. The pharmacokinetics and dosing of oral 4-methylumbelliferone for inhibition of hyaluronan synthesis in mice. Clin Exp Immunol. 2016;185(3):372–381. doi:10.1111/cei.12815

55. Hamzé R, Delangre E, Tolu S, et al. Type 2 Diabetes Mellitus and Alzheimer’s Disease: Shared Molecular Mechanisms and Potential Common Therapeutic Targets. Int J Mol Sci. 2022;23(23):15287. doi:10.3390/ijms232315287

56. O’Leary TP, Brown RE. Visuo-spatial learning and memory impairments in the 5xFAD mouse model of Alzheimer’s disease: Effects of age, sex, albinism, and motor impairments. Genes Brain Behav. 2022;21(4):e12794. doi:10.1111/gbb.12794

57. Silva A, Martínez MC. Spatial memory deficits in Alzheimer’s disease and their connection to cognitive maps’ formation by place cells and grid cells. Front Behav Neurosci. 2023;16:1082158. doi:10.3389/fnbeh.2022.1082158

58. Barrett GL, Bennie A, Trieu J, Ping S, Tsafoulis C. The chronology of age-related spatial learning impairment in two rat strains, as tested by the Barnes maze. Behav Neurosci. 2009;123(3):533–538. doi:10.1037/a0015063

59. Wilson E, Knudson W, Newell-Litwa K. Hyaluronan regulates synapse formation and function in developing neural networks. Sci Rep. 2020;10(1):16459. doi:10.1038/s41598-020-73177-y

60. Peters A, Sherman LS. Diverse Roles for Hyaluronan and Hyaluronan Receptors in the Developing and Adult Nervous System. Int J Mol Sci. 2020;21(17):5988. doi:10.3390/ijms21175988

61. Arranz AM, Perkins KL, Irie F, et al. Hyaluronan deficiency due to Has3 knock-out causes altered neuronal activity and seizures via reduction in brain extracellular space. J Neurosci Off J Soc Neurosci. 2014;34(18):6164–6176. doi:10.1523/JNEUROSCI.3458-13.2014

62. Foscarin S, Raha-Chowdhury R, Fawcett JW, Kwok JCF. Brain ageing changes proteoglycan sulfation, rendering perineuronal nets more inhibitory. Aging. 2017;9(6):1607–1622. doi:10.18632/aging.101256

63. Miyata S, Kitagawa H. Chondroitin 6-Sulfation Regulates Perineuronal Net Formation by Controlling the Stability of Aggrecan. Neural Plast. 2016;2016:1305801. doi:10.1155/2016/1305801

64. Yang S, Gigout S, Molinaro A, et al. Chondroitin 6-sulphate is required for neuroplasticity and memory in ageing. Mol Psychiatry. 2021;26(10):5658–5668. doi:10.1038/s41380-021-01208-9

65. Hendrickson AS, Francis KL, Kumar A, et al. Assessing translational applicability of perineuronal net dysfunction in Alzheimer’s disease across species. Front Neurosci. 2024;18. doi:10.3389/fnins.2024.1396101

66. Watanabe A, Hirayama S, Kominato I, et al. HAPLN2 forms aggregates and promotes microglial inflammation during brain aging in mice. PLOS Biol. 2025;23(8):e3003006. doi:10.1371/journal.pbio.3003006

67. Wilcock DM, Morgan D, Gordon MN, et al. Activation of matrix metalloproteinases following anti-Aβ immunotherapy; implications for microhemorrhage occurrence. J Neuroinflammation. 2011;8:115. doi:10.1186/1742-2094-8-115

