## Supplementary figures and images for "4-methylumbelliferone attenuates amyloid pathology and learning deficits in the APP/PS1 mouse model"

### Supplemental fig 1

#### Supplemental Figure 1

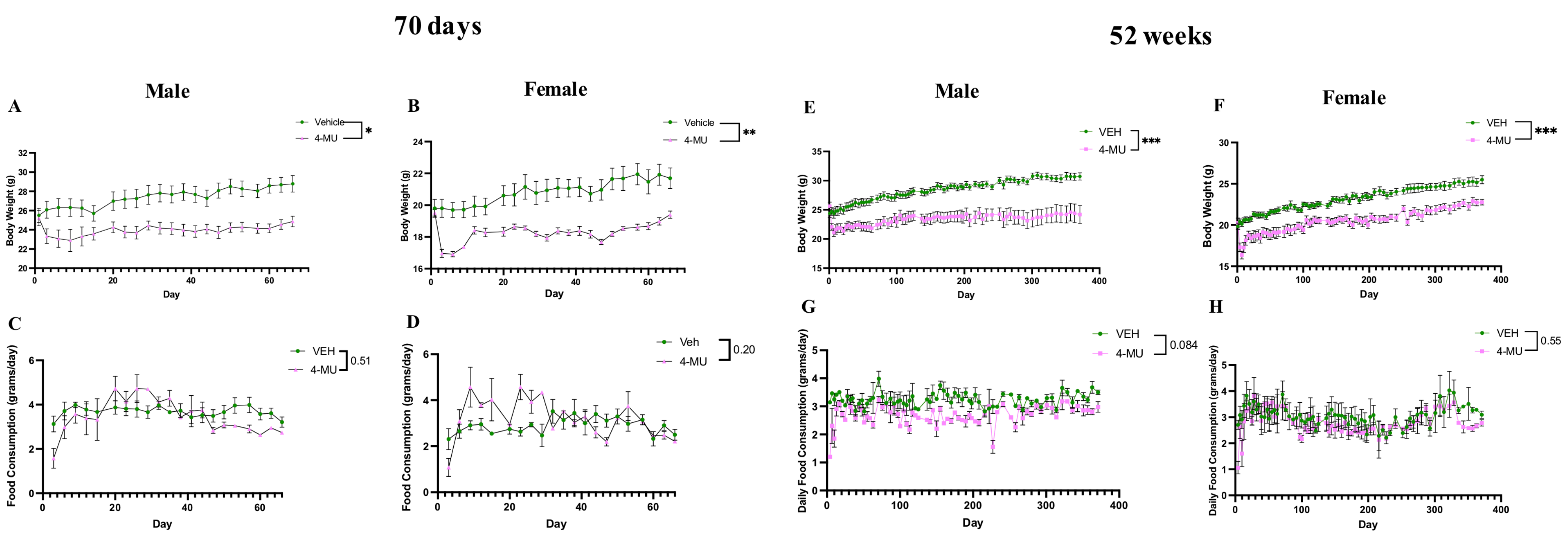

### Supplemental fig 2

#### Supplemental Figure 2

A

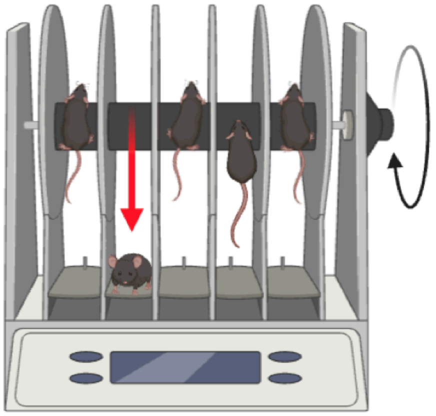

B

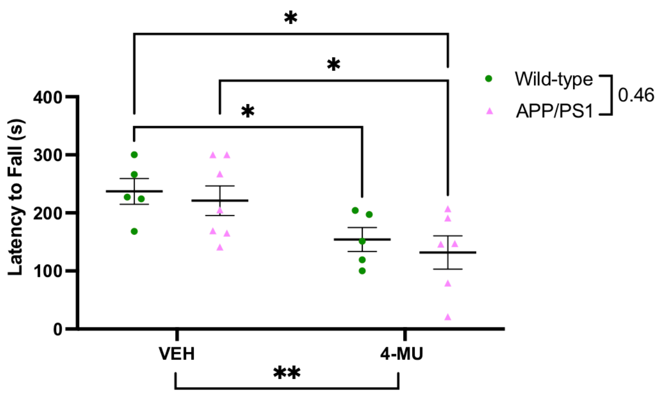


C

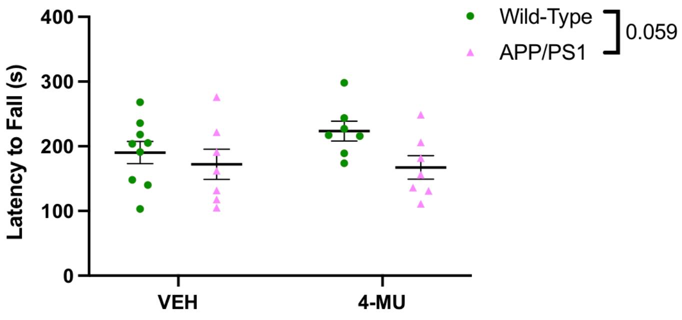

### Supplemental fig 3

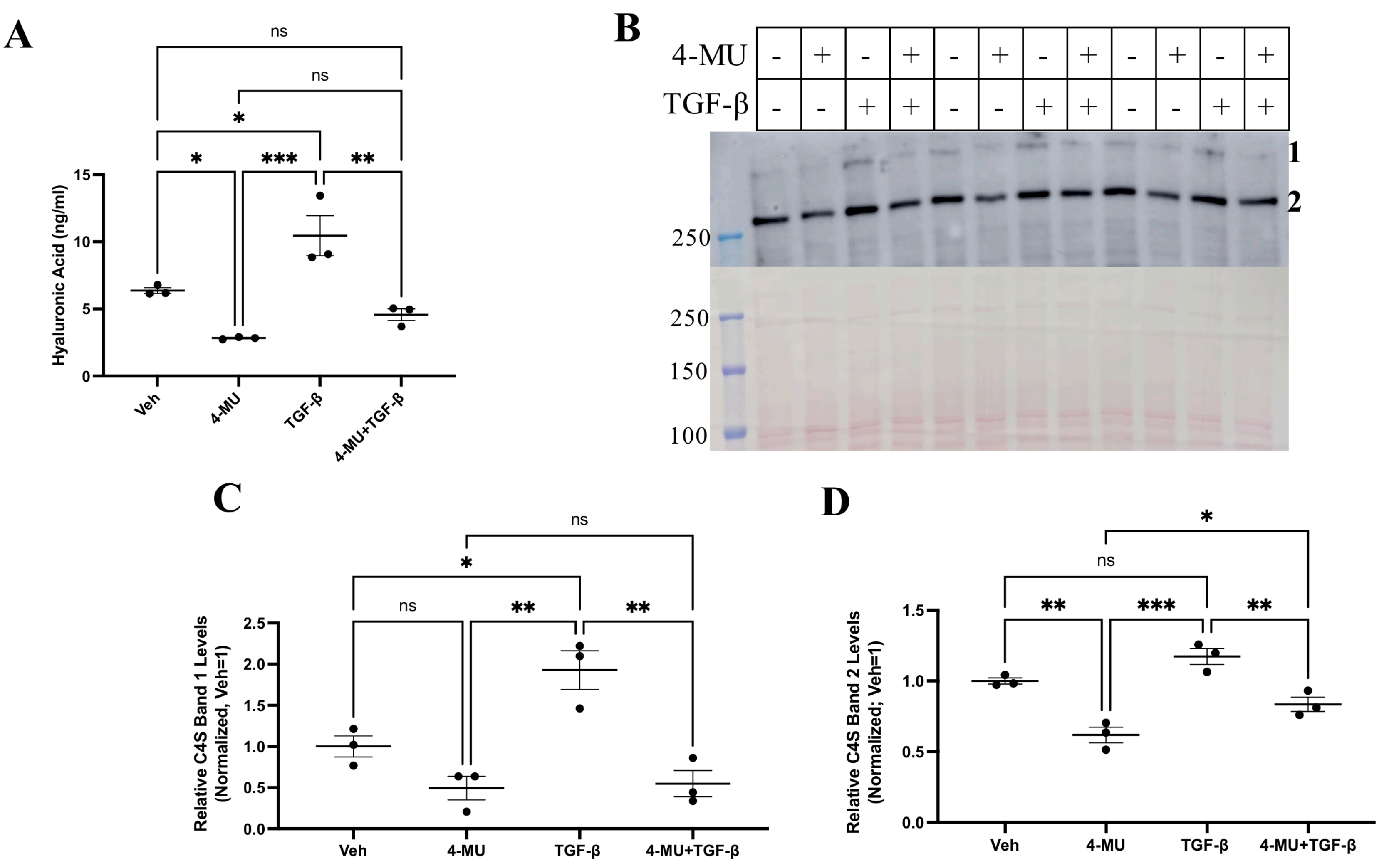
